# High Resolution *Salmonella* Subtyping by the MLSTnext Method

**DOI:** 10.1101/2023.03.10.532158

**Authors:** Zhihai Ma, Baback Gharizadeh, Steven Huang, Mo Jia, Florence Wu, Chunlin Wang

## Abstract

The food industry needs a straightforward, efficient, widely applicable and cost-efficient method to not only detect, but also determine serovar and potential sources of multiple strains in food and environment. While Whole Genome Sequencing (WGS) can generate complete genomic profile of food pathogens, it is a laborious, time-consuming, and expensive method that necessitates pure isolates. As a result, it is unsuitable for samples with complex background, limiting its widespread application by food industry. However, traditional multilocus sequence typing (MLST) approaches do not provide sufficient single nucleotide polymorphism (SNP) information to effectively track-and-trace sources of contamination. In contrast, ChapterDx MLSTnext-NGS (next-generation sequencing) *Salmonella* assay amplifies and sequences 47 polymorphic loci, evenly spanning the *Salmonella enterica* genome. As demonstrated in this study, the ChapterDx MLSTnext-NGS *Salmonella* can identify serovar with high resolution, distinguishing between various strains of the same serovar and resolve co-presence of different strains in a single sample. Additionally, this assay can analyze up to thousand samples in a single sequencing run within 22 hours, making it a highly efficient and scalable method for the food industry. Moreover, the cost per sample of the ChapterDx MLSTnext-NGS *Salmonella* assay is comparable to that of quantitative Polymerase Chain Reaction (qPCR), making it an affordable option. The assay uses ChapterDx amplification technology, allowing for the amplification of all 47 loci in a single-tube, single-step PCR reaction. This makes it one of the simplest NGS applications available. In summary, ChapterDx MLSTnext-NGS *Salmonella* assay can be implemented in many diagnostic laboratories to address increasingly complex food safety issues.

## Introduction

Globalization has increased the complexity of the food supply chain as more than ever; companies are relying on ingredient and raw materials suppliers from more regions of the world. This in turn has led to a potential increase in risk associated with the food as the raw materials may not be produced under the same hygienic standards as those produced domestically. This has resulted in changes in food safety regulations, for both domestic and foreign food suppliers, requiring them to identify all potential hazards associated with the food and implement measures to prevent these risks from becoming a risk to the public health. Despite these measures, food safety recalls due to product contamination as well as outbreaks, especially those concerning pathogenic microorganisms, continue to occur at significant cost to the public and food manufacturers. Among the preventive measures identified is the need for better technologies that can allow various players within the food supply chain to proactively detect, identify, and remove contaminated foods.

Bacterial subtyping is an important measure in preventing foodborne illness outbreaks, as it allows public health officials to quickly identify the source of contamination and take appropriate action to prevent further spread. There are several methods of bacterial subtyping that are currently being used. Serotyping^1^ is a phenotypic typing method that involves identifying the type of antigens on the surface of the bacterial cell. This method can be used to differentiate between different strains of a species, but it is less discriminatory than molecular methods. Pulsed-field gel electrophoresis (PFGE)^2^ is a molecular typing method that uses restriction enzymes to digest bacterial DNA into large fragments, which are then separated by size using gel electrophoresis. The resulting banding patterns can be used to compare bacterial strains and identify relatedness. MLST^3^ is a molecular typing method that involves sequencing specific genes within the bacterial genome. The resulting sequences can be compared with the sequences from other relevant strains to identify their relatedness and to infer their evolutionary relationships. WGS^4^ is a molecular typing method that sequences the entire bacterial genome. This method provides the highest level of resolution and can be used to identify SNPs and other genetic variations within the pathogen. WGS can provide the highest level of specificity and sensitivity in identifying bacterial subtypes and tracking the spread of outbreaks.

Traditional MLST identifies bacterial strains based on the sequence variation of several housekeeping genes. For instance, *Salmonella* MLST sequences seven housekeeping genes of *Salmonella*: *chorismate synthase (aroC), β sliding clamp of DNA polymerase III (dnaN), uroporphyrinogen-III synthase (hemD), histidinol dehydrogenase (hisD), phosphoribosylaminoimidazole carboxylase catalytic subunit (purE), 2-oxoglutarate dehydrogenase E1 component (sucA) and homoserine dehydrogenase (thrA)*^5^. These housekeeping genes are highly conserved and only slowly accumulate site changes^6^, so *Salmonella* MLST often lacks the discriminatory power to differentiate strains of the same serovar or closely-related serovar, limiting its use in determining relatedness of the isolates and in source tracking during investigations. In order to increase the discriminating power of MLST, a new strategy named MLSTnext has been developed by Chapter Diagnostics Inc. that can analyze sequence variations of several folds more regions than the traditional MLST, allowing it to detect sequence variations across much larger regions of the bacterial genome. MLSTnext examines regions beyond just housekeeping genes. These regions are more heterogeneous, meaning they have greater diversity, and some loci may be absent in certain strains. This increased diversity allows for better differentiation among strains of the same bacterial species, leading to a more accurate analysis of genetic relatedness. Here we introduce ChapterDx MLSTnext-NGS *Salmonella* assay, a user-friendly NGS-based technology that can subtype *Salmonella* down to the sub-strains level. This innovative assay offers high accuracy in serotype identification, enabling the detection of sub-strains and resolution of multiple serovars in the same sample.

## Method and Material

### Sample

For this study, 12 ATCC *Salmonella* isolates and three other isolates were used (Table 1). Stocks of the bacterial strains preserved in glycerol were recovered from −80°C stock storage. Isolates were streaked on Tryptic Soy Agar with 5% Sheep Blood plates and incubated at 35°C overnight. One to three colonies were picked, and their DNA were extracted using the QIAamp Mini DNA kit following the manufacturer’s protocol. *Salmonella* isolates were propagated, and DNA was extracted at a collaborating laboratory (AEMTEK Inc., Fremont, CA).

**Table 1.** Information about samples used in this study

| No | Species and subspecies | Serovar | Source | Origin |
| --- | --- | --- | --- | --- |
| 1 | *S. enterica* subsp. *enterica* | Abaetetuba | ATCC 35640 | Creek water |
| 2 | *S. enterica* subsp. *enterica* | Anatum | ATCC 9270 | Pork liver |
| 3 | *S. enterica* subsp. *enterica* | Bispebjerg | ATCC 9842 | Unknown |
| 4 | *S. enterica* subsp. *enterica* | Diarizoniae | ATCC 12325 | Unknown |
| 5 | *S. enterica* subsp. *enterica* | Infantis | BAA-1675 | Unknown |
| 6 | *S. enterica* subsp. *enterica* | Mbandaka | ATCC 51958 | Unknown |
| 7 | *S. enterica* subsp. *enterica* | Montevideo | ATCC 8387 | Unknown |
| 8 | *S. enterica* subsp. *enterica* | Poona | NCTC: 4840 | Infant enteritis |
| 9 | *S. enterica* subsp. *enterica* | Seftenberg | ATCC 8400 | Unknown |
| 10 | *S. enterica* subsp. *enterica* | Tennessee | ATCC 10722 | Unknown |
| 11 | *S. enterica* subsp. *enterica* | Typhi | ATCC 6539 | Unknown |
| 12 | *S. enterica* subsp. *enterica* | Typhimurium | ATCC 14028 GFP | Chicken tissue |
| 13 | *S. enterica* subsp. *enterica* | Unknown | Unknown | Unknown |
| 14 | *S. enterica* subsp. *enterica* | Typhimurium | Unknown | Unknown |
| 15 | *S. enterica* subsp. *enterica* | Typhimurium | Unknown | Unknown |

### Assay design

ChapterDx MLSTnext-NGS *Salmonella* assay examines 47 polymorphic loci from *Salmonella enterica* genome. The 47 loci include seven loci for housekeeping genes (*aro*C, *dna*N, *hem*D, *his*D, *pur*E, *suc*A, and *thr*A) used in traditional MLST assays. Although the assay is designed based on *S. enterica* genome, more than 20 loci could get amplified and sequenced for *Salmonella bongori* (data not shown).

### PCR

One-step multiplex PCR was performed in 15 μl final volume in a 96-well plate on a Veriti thermocycler (ThermoFisher, CA, USA). The PCR reaction consisted of target-specific primers, barcoded universal primers, sample DNA, DNA polymerase, dNTPs and PCR buffer. The PCR conditions were according to the ChapterDx MLSTnext-NGS *Salmonella* assay instructions.

### Library preparation and next-generation sequencing

After PCR, the amplified products were quenched and pooled into a 1.5 ml tube (or a 15 ml tube depending on the sample size). A portion of the pooled amplicons was purified with SPRIbeads (Beckman Coulter, CA, USA) following the ChapterDx MLSTnext-NGS *Salmonella* assay instructions. The purified pooled amplicons concentration was measured on a Qubit 3 fluorometer (ThermoFisher, CA, USA), and the concentration was adjusted for sequencing according to Illumina library preparation instructions. The library was sequenced on an Illumina MiniSeq platform using an Illumina Mid-Output sequencing kit (Illumina, CA, USA). The workflow is depicted in Figure 1.

**Figure 1.**
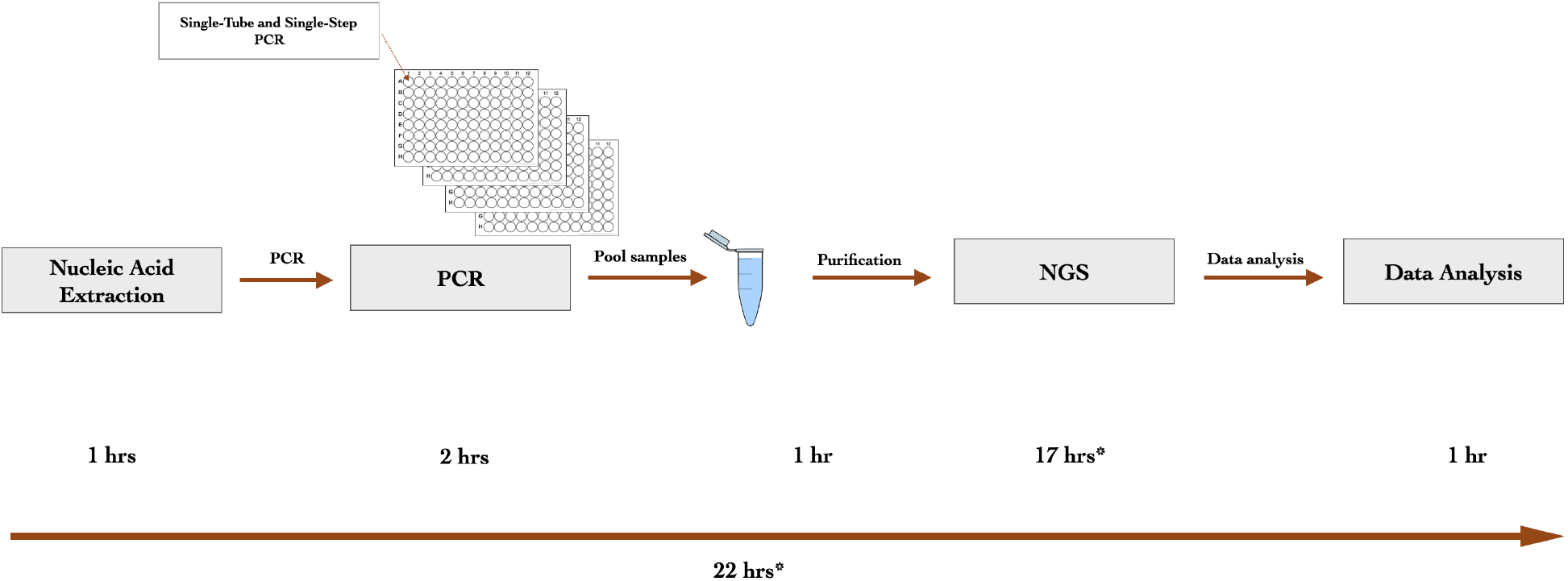
A schematic illustration of ChapterDx MLSTnext-NGS *Salmonella* assay workflow. *The overall time depends on the sequencing kit. It takes about 17 hours to complete sequencing with Illumina Miniseq Mid-output kit.

### Sequence data analysis software and interpretation

Decoding: sequencing reads (FASTQ format) for forward and reverse reads and forward and reverse indexing reads were input in ChapterDx Analysis Software. The software assigns sequencing reads for a sample based on the sequence of both forward and reverse index reads with no mismatch; Mapping: sequencing reads are mapped onto reference sequences using the Smith-Waterman algorithm^7^ with options as nucleotide match reward is 1, nucleic mismatch penalty is - 3, cost to open a gap is 5, and cost to extend a gap is 2. Only the alignment of best match is kept for each sequencing read if the alignment score is beyond 60. Subtyping: 1) find the genome (X) with most mapped loci; 2) exit if the number of mapped loci is less than pre-set cutoff; 3) output genome (X) and remove all mapped reads to genome(X); 4) repeat step 1-3. For each locus, paired (one forward and one reverse) consensus sequences was generated. Pairwise distance was calculated as the number of allele difference between two strains. Neighbor-joining tree was then constructed based on the resultant distance matrix.

## Result

### Distance between Samples

Table 2 lists the distances between each pair of tested samples. Pairwise distance was calculated as the number of allele difference between two strains, which is the same as traditional MLST^11^. In traditional MLST the number of nucleotide differences between alleles is ignored and sequences are given different allele numbers whether they differ at a single nucleotide site or at many sites. The rationale is that a single genetic event resulting in a new allele can occur by a point mutation (altering only a single nucleotide site), or by a recombinational replacement (that will often change multiple sites). From Table 2, isolates of different serovars differ more than 30 loci, suggesting that loci examined by ChapterDx assay are heterogenous. Sample 12 and 15 are both *S. typhimurium* serovar. However, alleles at four loci are different for these two samples of the same serovar. These four loci do not overlap any of the seven housekeeping genes (*aro*C, *dna*N, *hem*D, *his*D, *pur*E, *suc*A, and *thr*A) used in traditional MLST. Thus, traditional MLST would have unlikely discriminated them.

**Table 2.** Distance matrix for tested samples.

|  | Abaetetuba(1<sup>*</sup>) | Anatum(2) | Bispejerg(3) | Diarizoniae(4) | Infantis(5) | Mbandaka(6) | Montevideo(7) | Poona(8) | Senftenberg(9) | Tennessee(10) | Typhi(11) | Typhimurium(12) | Enteritidis(13) | Muenster(14) | Typhimurium(15) |
| --- | --- | --- | --- | --- | --- | --- | --- | --- | --- | --- | --- | --- | --- | --- | --- |
| Abaetetuba(1) | 0<sup>#</sup> | 44 | 44 | 47 | 36 | 41 | 30 | 33 | 41 | 44 | 45 | 46 | 45 | 38 | 45 |
| Anatum(2) | 44 | 0 | 41 | 47 | 43 | 42 | 41 | 42 | 37 | 37 | 44 | 40 | 39 | 44 | 40 |
| Bispejerg(3) | 44 | 41 | 0 | 47 | 46 | 42 | 44 | 46 | 45 | 44 | 46 | 40 | 39 | 46 | 40 |
| Diarizoniae(4) | 47 | 47 | 47 | 0 | 47 | 47 | 47 | 47 | 47 | 47 | 47 | 47 | 47 | 47 | 47 |
| Infantis(5) | 36 | 43 | 46 | 47 | 0 | 45 | 34 | 40 | 42 | 43 | 46 | 45 | 46 | 37 | 44 |
| Mbandaka(6) | 41 | 42 | 42 | 47 | 45 | 0 | 46 | 46 | 44 | 44 | 45 | 42 | 43 | 44 | 43 |
| Montevideo(7) | 30 | 41 | 44 | 47 | 34 | 46 | 0 | 38 | 40 | 43 | 46 | 45 | 46 | 36 | 44 |
| Poona(8) | 33 | 42 | 46 | 47 | 40 | 46 | 38 | 0 | 45 | 44 | 45 | 45 | 44 | 35 | 45 |
| Senftenberg(9) | 41 | 37 | 45 | 47 | 42 | 44 | 40 | 45 | 0 | 38 | 45 | 40 | 45 | 44 | 40 |
| Tennessee(10) | 44 | 37 | 44 | 47 | 43 | 44 | 43 | 44 | 38 | 0 | 46 | 44 | 43 | 46 | 44 |
| Typhi(11) | 45 | 44 | 46 | 47 | 46 | 45 | 46 | 45 | 45 | 46 | 0 | 44 | 44 | 45 | 44 |
| Typhimurium(12) | 46 | 40 | 40 | 47 | 45 | 42 | 45 | 45 | 40 | 44 | 44 | 0 | 40 | 45 | 4 |
| Enteritidis(13) | 45 | 39 | 39 | 47 | 46 | 43 | 46 | 44 | 45 | 43 | 44 | 40 | 0 | 45 | 41 |
| Muenster(14) | 38 | 44 | 46 | 47 | 37 | 44 | 36 | 35 | 44 | 46 | 45 | 45 | 45 | 0 | 45 |
| Typhimurium(15) | 45 | 40 | 40 | 47 | 44 | 43 | 44 | 45 | 40 | 44 | 44 | 4 | 41 | 45 | 0 |

\*The number inside parenthesis is sample Id (see Table 1 and 3). <sup>#</sup> indicates the number of loci different between two samples.

### Subtyping

The serotype calling algorithm infers the serovar from the best-matched genome assembly. Often, the best-matched genome assemblies are annotated with serovar information. However, sometimes, the best-matched genome assemblies lack serovar information. In order to fill the missing serovar for genome assemblies, several difference algorithms^8,9^ have been tested to estimate the evolutionary distances among *Salmonella* assemblies downloaded from the NCBI database. Eventually, the program andi^8^ was chosen to calculate the distance between two Salmonella genome assemblies because the resultant distance matrix reflected the relationships among *Salmonella* genomes. Based on the calculated distance matrix, many *Salmonella* genome assemblies without serovar information were grouped together with those with serovar information using the single-linkage clustering algorithm^10^. Table 3 lists the NCBI accessions for best-matched genome assemblies and their corresponding inferred serovars. There is no serovar information for sample 13 that ChapterDx assay identified serotype Enteritidis. Sample 14 was labeled as *S. typhimurium* originally, but our assay identified it as *S. muenster* with best-matched genome accession CP019198.1. The neighbor-joining tree (Figure 2) based on allele difference shows that sample 14 is separated from both sample 12 (*S. typhimurium*) and sample 15 (*S. typhimurium*) with 45 loci difference (Table 2), suggesting that sample 14 is less likely to be *S. typhimurium*.

**Table 3.**
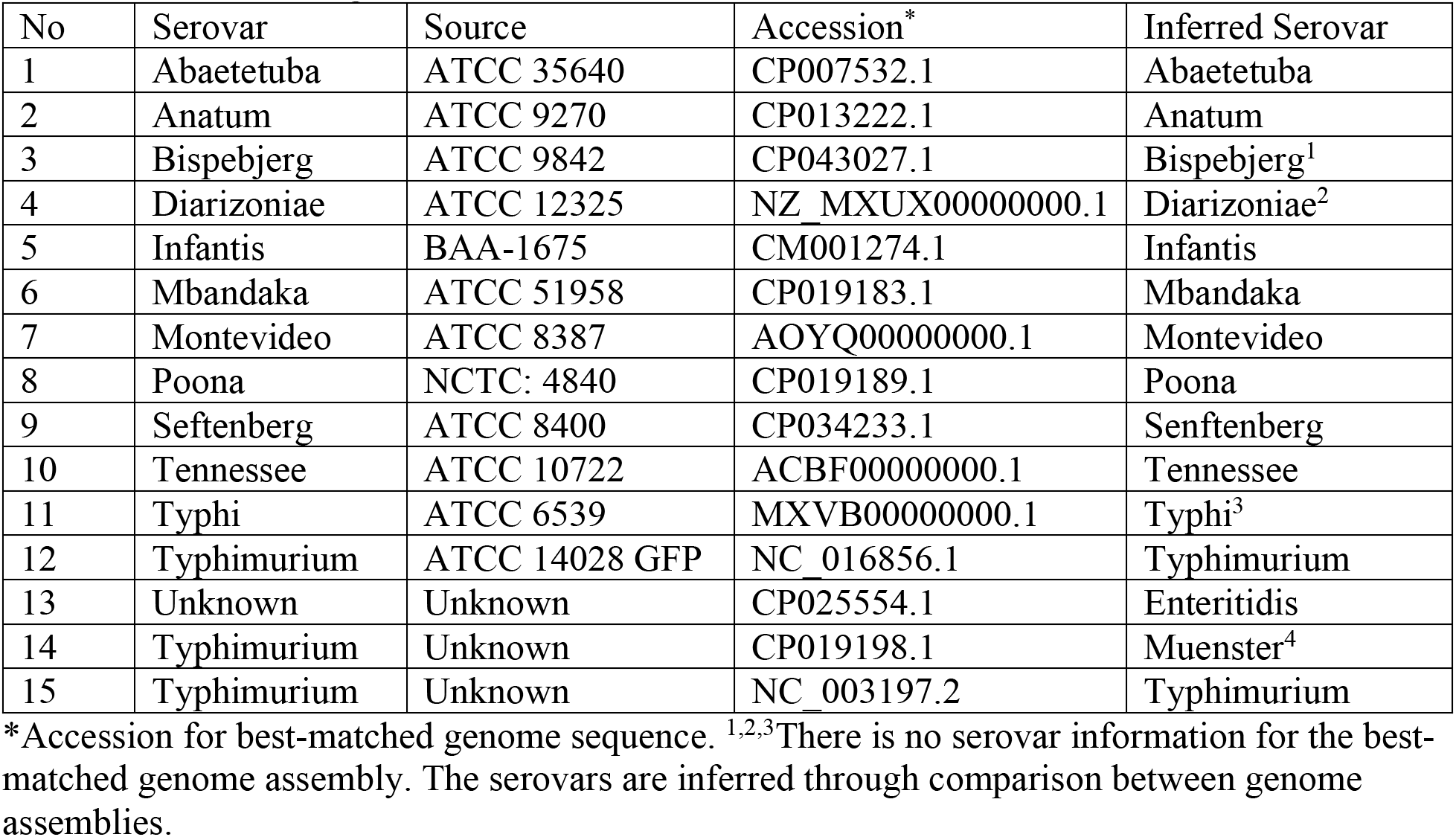
Best-matched genome accession and inferred serovar.

**Figure 2.**
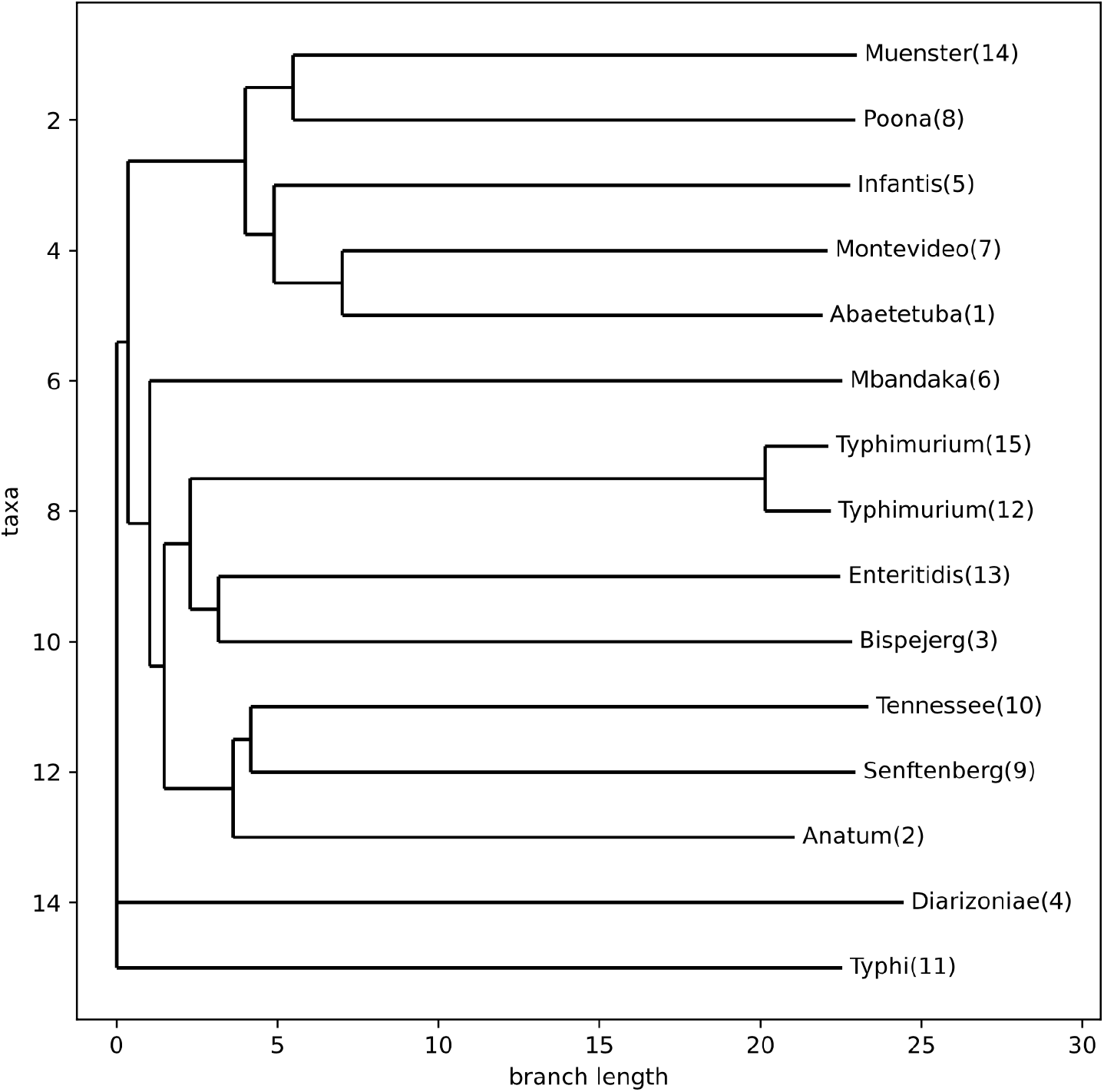
Neighbor-joining tree for tested samples. The number inside parenthesis is sample Id (see Table 1 and 3).

### Detecting co-presence of multiple strains in the same sample

It has been reported that the same food or environmental samples could be contaminated with different *Salmonella* strains simultaneously^12^. As such multiple serotype *Salmonella* outbreaks^13–15^ might occur more frequently than recognized. Identification of all *Salmonella* serotypes involved in an outbreak might help implicate the outbreak source, define the scope of the outbreak, and determine the selection of appropriate control measures. To evaluate ChapterDx assay’s capability to detect co-presence of multiple strains in the same samples, DNA from multiple samples were mixed to mimic samples contaminated with multiple strains. The experiment was performed in mixtures of 3, 6 and 9 serovars, where 1) three tubes contained mixture of three serovars each, 2) three tubes contained mixture of six serovars each, and 3) two tubes contained mixtures of nine serovars each. The two mixture tubes with nine serovars were identical. The results show that ChapterDX MLSTnext assay could identify and differentiate all serovar by ChapterDx automatic data analysis software from all mixtures, suggesting this assay’s excellent capability to detect co-presence of multiple strains in the same sample (Table 4).

**Table 4.** Mixture sample information and result

| Sample Id | Mixture | Result |
| --- | --- | --- |
| Mix3_1 | Sample 1, 2, 3 | Match |
| Mix3_2 | Sample 5, 6, 7 | Match |
| Mix3_3 | Sample 8, 9, 10 | Match |
| Mix6_1 | Sample 1, 2, 3, 5, 6, 7 | Match |
| Mix6_2 | Sample 1, 2, 3, 8, 9, 10 | Match |
| Mix6_3 | Sample 5, 6, 7, 8, 9, 10 | Match |
| Mix9_1 | Sample 1, 2, 3, 5, 6, 7, 8, 9 10 | Match |
| Mix9_2 (repeat) | Sample 1, 2, 3, 5, 6, 7, 8, 9 10 | Match |

## Discussion

WGS analyzes the entire genome of a bacterial strain and provides a comprehensive overview and details of the genetic content of the strain. WGS is a powerful tool for food safety-related investigations, but it has some limitations. First, WGS generates a large amount of data, which can be challenging to analyze. Analysis requires powerful computing resources and specialized bioinformatics expertise. Second, WGS is time and labor intensive and expensive, which can be a barrier to its use in some settings and thus, is never used as a routine test. Third, the lack of standardization in data analysis and interpreting the results can be challenging. Genetic variation within a bacterial species can be complex, and it can be difficult to determine the significance of differences between strains. Lastly, WGS requires pure colony isolate, which limits its usage on complex samples.

Traditional MLST classifies and compares bacterial strains based on the nucleotide sequences of several gene loci. The method involves PCR amplification and sequencing internal fragments of 6-8 housekeeping genes, which are genes that encode for essential metabolic functions and are conserved among bacterial species. Although traditional MLST can differentiate between bacterial strains based on sequence variations at targeted loci, it may not be able to resolve closely related strains.

MLSTnext occupies a middle ground between traditional MLST and WGS in terms of its resolution and complexity. MLSTnext characterizes several folds more loci than traditional MLST, and the extra loci are more diverse than those housekeeping genes examined by traditional MLST. MLSTnext-based ChapterDX NGS *Salmonella* assay characterizes 47 loci, as compared to 7 housekeeping genes characterized by traditional MLST methods, thus significantly increasing its discriminatory power. This enables accurate serotype identification, distinguishes strains of the same serovar, and detects co-presence of multiple strains in the same sample. To date, this is the only known method capable of detecting and characterizing multiple strains in the same sample, thus, reducing the need to run multiple assay that can be very expensive.

Prompt identification of *Salmonella* serovars from diagnostic samples is critical to enable timely implementation of appropriate management decisions. Although identification of *Salmonella* at the serogroup level can be rapidly achieved at most diagnostic laboratories, final serovar identification often takes several weeks because of the limited number of reference laboratories performing the complex task of serotyping. While MLSTnext characterizes many more loci than traditional MLST, with ChapterDX proprietary amplification strategy^16^, all 47 loci can be amplified with single-tube, single-step PCR (Figure 1). With a next-generation sequencing (NGS) platform such as Illumina sequencing machine, hundreds to thousands of samples can be analyzed in the same sequencing run within a 22-hour period. Such a high-level multiplexing can significantly lower the cost per sample, making it comparable to qPCR. This assay comes with customized software to analyze the sequence data and to generate report. No bioinformatics expertise is needed to carry out data analysis. Therefore, this assay can be implemented in many diagnostic laboratories.

MLSTnext is a generic strategy. It has been applied to subtype not only bacteria such as *Listeria*, *Legionella*, Shiga toxin-producing *E. coli* (STEC), parasites such as *Cyclospora*, but also viruses such as Hepatitis A virus. Coupled with the ChapterDx amplification technology^16^ and NGS, MLSTnext could be a straightforward, effective, widely applicable and cost-efficient method to address increasingly complex food safety issues.

